# Concomitant acetylation and loading of H2A.Z by NuA4/TIP60 regulate target gene transcription

**DOI:** 10.1101/2025.09.22.677705

**Authors:** Amel Mameri, Catherine Lachance, Claudia Cattoglio, Stéphanie Bianco, Jonathan Humbert, Charles Joly-Beauparlant, Emeric Texeraud, Arul Banerjea, Anahita Lashgari, Jean-Philippe Lambert, Samer M. I. Hussein, Arnaud Droit, Jacques Côté

## Abstract

The human NuA4/TIP60 complex is a multi-subunit, dual enzymatic epigenetic factor and gene regulator. It bears histone acetyltransferase (HAT) activity towards the canonical histones H2A and H4 and the histone variant H2A.Z, a function that has been linked to gene activation. It also acts as a chromatin remodeling enzyme through ATP-dependent exchange of nucleosomal H2A-H2B dimers with H2A.Z-H2B, leading to incorporation of H2A.Z into chromatin at gene regulatory elements. NuA4/TIP60 is unique in merging two enzymatic activities targeting H2A.Z. Both NuA4/TIP60-dependent H2A.Z acetylation and remodeling have been linked to several physiological functions and pathologies, but studies have only focused on either one or the other enzymatic activity, and insights on functional coordination between them are lacking. Here, we leverage our EP400 rapid depletion system to explore and untangle the intricate links between H2A.Z acetylation by Tip60 (the HAT subunit) and loading on chromatin by EP400 (the remodeling subunit) through functional genomic and biochemical analyses. Our data support a mechanism in which H2A.Z is first pre-acetylated to allow for H2A.Zac-H2B dimer association with the complex before incorporation into chromatin, particularly at gene promoters to positively regulate transcription. As both H2A.Z-targeted enzymatic functions of NuA4/TIP60 have been linked to disease, albeit separately, our findings hold important implications for therapeutic intervention, where combinatorial targeting is a promising avenue.

## INTRODUCTION

DNA-templated reactions in eukaryotes, including gene transcription, are controlled by factors which integrate and translate cellular signals into structural rearrangements of chromatin through enzymatic activities targeting the nucleosomes and altering DNA accessibility. For example, the histones composing the nucleosomes are subject to covalent modifications by chromatin modifying enzymes. Furthermore, replacement of canonical, replication-dependent histones with their non-canonical, replication-independent variants, carried out by ATP-dependent chromatin remodeling enzymes, is another mechanism regulating nucleosome structural properties. Both chromatin modifying enzymes and remodeling enzymes mainly act as multiprotein complexes, sometimes merging different enzymatic activities. They often contain “reader” modules that are able to recognize specific histone modifications, facilitating recruitment of the complex to specific genomic sites and allowing for functional crosstalk between the chromatin modulating machineries (Millan-Zambrano et al., 2022; Venkatesh & Workman, 2015).

H2A.Z is a variant of the canonical histone H2A and is essential in mammals. It is incorporated throughout the cell cycle into various regions across the genome, including euchromatin which contains actively transcribed genes, facultative heterochromatin containing repressed genes, and constitutive heterochromatin particularly in pericentric regions where it plays a role in chromosome segregation. H2A.Z has been shown to regulate all three major DNA-templated processes, with roles in DNA break repair, DNA replication, and gene transcription (Colino-Sanguino et al., 2022; Giaimo et al., 2019). Its role in transcriptional regulation is context- dependent, where post-translational modifications (PTMs) of H2A.Z N- and C-terminal tails significantly impact and define the transcriptional output resulting from its incorporation into gene promoters and enhancers. Acetylation of lysine residues on the N-terminal tail of H2A.Z activates gene transcription, whereas ubiquitination of C-terminal lysines represses it (Giaimo et al., 2019). Differences between H2A.Z isoforms (H2A.Z.1, H2A.Z.2.1 and H2A.Z.2.2 in mammals) also contribute to the complexity of its function in transcription with their different interactomes and variable effects on nucleosome stability (Giaimo et al., 2019; Lamaa et al., 2020).

Two human ATP-dependent remodelers have been described to perform the H2A-H2B to H2A.Z-H2B dimer exchange, incorporating H2A.Z into chromatin: SRCAP, the remodeler subunit found in the SRCAP complex (Bowman et al., 2011; Ruhl et al., 2006; Slupianek et al., 2010; Wong et al., 2007; Yang et al., 2012; Yu et al., 2024), and its paralog EP400, which is part of the NuA4/TIP60 complex (Bellucci et al., 2013; Bowman et al., 2011; Gevry et al., 2007; Lashgari et al., 2017; Sudarshan et al., 2022; Xu et al., 2012). Studies in mammals mainly describe the SRCAP complex as a transcription co-factor (Bowman et al., 2011; Patty & Hainer, 2024; Slupianek et al., 2010; Wong et al., 2007; Yang et al., 2012; Yu et al., 2024), with roles in cell cycle regulation (Chen et al., 2023; Jostes et al., 2024; Sun et al., 2022), cell fate determination (Cuadrado et al., 2010; Ye et al., 2017; Zhao et al., 2019), heart development and function (Shi et al., 2022; Xu et al., 2021), benign and cancerous tumorigenesis (Berta et al., 2021; Jostes et al., 2024; Pugacheva et al., 2024; Zhang et al., 2024), and neurodevelopmental disorders (Ding et al., 2024; Greenberg et al., 2019). While the remodeling function of mammalian EP400 has received less attention than that of SRCAP, it has also been mostly linked to transcriptional regulation via H2A.Z exchange at *cis*-regulatory elements (Bellucci et al., 2013; Bowman et al., 2011; Couture et al., 2012; Frob et al., 2019; Gevry et al., 2007; Jostes et al., 2024; Pradhan et al., 2016; Sudarshan et al., 2022; Sun et al., 2023; Yang et al., 2024) and to a lesser extent to DNA damage repair through incorporation of the variant at break sites (Begum et al., 2021; Xu et al., 2012). Similar to SRCAP, gene regulation through EP400-mediated H2A.Z deposition is involved in cell cycle regulation (Bellucci et al., 2013; Gevry et al., 2007; Jostes et al., 2024; Yang et al., 2024), cell differentiation (Couture et al., 2012), cancer (Jostes et al., 2024; Sun et al., 2023), neurodevelopment and neuronal function (Frob et al., 2019). Moreover, the SRCAP and the NuA4/TIP60 complexes share several subunits which have also been implicated in H2A.Z exchange (Gallant-Behm et al., 2012; Hsu, Shi, et al., 2018; Hsu, Zhao, et al., 2018; Jostes et al., 2024; Kikuchi et al., 2023; Latrick et al., 2016; Tartour et al., 2022) and linked to transcriptional regulation (Gallant-Behm et al., 2012; Hsu, Shi, et al., 2018; Hsu, Zhao, et al., 2018; Jostes et al., 2024; Kikuchi et al., 2023; Tartour et al., 2022), cancer cell cycle progression (Hsu, Shi, et al., 2018; Jostes et al., 2024), embryonic stem cell maintenance (Hsu, Zhao, et al., 2018) and regulation of the circadian rhythm (Tartour et al., 2022).

The existence of two paralogous remodelers of H2A.Z with seemingly overlapping molecular and physiological functions raises the question of what the redundant and specific roles of EP400 and SRCAP are within the mammalian cell. In addition to its remodeling activity, the human NuA4/TIP60 complex was first characterized as an acetyltransferase for the canonical histones H2A and H4 (Doyon et al., 2004), an enzymatic activity through which it acts as a gene co-activator controlling essential cellular and physiological processes (Devoucoux, Fort, et al., 2022; Doyon et al., 2004; Humbert et al., 2020; Mameri & Cote, 2023; Sudarshan et al., 2022). Our past work on the budding yeast NuA4 complex (Altaf et al., 2010), and more recently on the human complex (Yang et al., 2024), provides evidence that acetylation of nucleosomal H2A and H4, mediated by the acetyltransferase subunit Tip60, enhances H2A.Z incorporation by EP400, suggesting that the remodeling function of the NuA4/TIP60 complex is coordinated with its acetyltransferase function, a mechanistic distinction setting it apart from SRCAP. Importantly, NuA4/TIP60 is the major acetyltransferase targeting H2A.Z in mammals (Colino-Sanguino et al., 2019; Dalvai et al., 2013; Giaimo et al., 2018; Ito et al., 2018; Janas et al., 2022; Jostes et al., 2024; Numata et al., 2020; Procida et al., 2021; Sporrij et al., 2023; Wichmann et al., 2022; Yamagata et al., 2021), where acetylation of chromatin-associated H2A.Z shows near complete dependence on Tip60 activity (Janas et al., 2022; Wichmann et al., 2022). In line with its role as a gene co- activator, NuA4/TIP60 acetylates H2A.Z on promoters and enhancers (Dalvai et al., 2013; Giaimo et al., 2018; Ito et al., 2018; Janas et al., 2022; Numata et al., 2020; Procida et al., 2021; Sporrij et al., 2023; Wichmann et al., 2022; Yamagata et al., 2021) to positively regulate transcription of genes involved in cell cycle progression (Dalvai et al., 2013; Wichmann et al., 2022), cell fate, including hematopoietic and neuronal cell fate (Janas et al., 2022; Numata et al., 2020; Sporrij et al., 2023), and cancer (Ito et al., 2018; Yamagata et al., 2021). Thus, H2A.Z is a substrate for the two enzymes within NuA4/TIP60, raising the possibility of simultaneous H2A.Z exchange and acetylation as a mechanism of gene activation that is specific to the NuA4/TIP60 complex, underlying its role in key cellular functions and pathologies.

To test this hypothesis, we combine biochemical analyses with functional genomic and transcriptomic analyses carried out in cell lines which allow acute EP400 depletion to interrogate the relationship between H2A.Z remodeling by EP400 and its acetylation by Tip60. We find strong evidence for EP400-mediated incorporation of pre-acetylated H2A.Z into chromatin to activate transcription at NuA4/TIP60 target genes, particularly those involved in cell proliferation, cell fate determination, development, and cancer. Given the physio-pathological relevance of both H2A.Z exchange and acetylation by NuA4/TIP60, the mechanistic insights provided by our study, intricately linking the two enzymatic activities targeting H2A.Z, can serve as a framework for therapeutic strategies.

## RESULTS

### EP400 remodeling activity is crucial for maintaining acetylated H2A.Z on chromatin

To investigate the functional and mechanistic links between remodeling and acetylation of H2A.Z by the human NuA4/TIP60 complex, we used K562 cell lines recently established by our group with the *EP400* gene homozygously tagged (EP400-FKBP12^F36V^-2xHA, referred to as “EP400- dTAG” from hereon) to allow for acute depletion of the endogenous EP400 protein upon treatment with the dTAG drug (Yang et al., 2024). Bulk analysis of chromatin in three different clones through Western blots of histones found in the chromatin enriched extracts (CEEs) of untreated (0h) or dTAG-treated cells at two timepoints (6h, 24h) shows a marked reduction in acetylated H2A.Z (H2A.Zac), a more subtle reduction in acetylated H4 (H4ac), and no change in total H2A.Z upon acute knock-out of EP400 (Figures 1A and S1A-B). The absence of observable change in total chromatin-bound H2A.Z suggests that only a small, undetectable fraction of H2A.Z found in the genome is actively remodelled by EP400, with the remaining sites either depending on SRCAP-mediated H2A.Z incorporation or having low turnover rates that are unaffected by short-term remodeler depletion, likely consisting of heterochromatin sites. Since EP400 acts as a major scaffold within the NuA4/TIP60 complex (Yang et al., 2024), the drop in histone acetylation following EP400 depletion is potentially attributable to loss of histone acetyltransferase (HAT) activity due to complex disintegration. The pronounced reduction in H2A.Zac is in agreement with the literature describing NuA4/TIP60 as the major HAT for H2A.Z (Colino-Sanguino et al., 2019; Dalvai et al., 2013; Giaimo et al., 2018; Ito et al., 2018; Janas et al., 2022; Jostes et al., 2024; Numata et al., 2020; Procida et al., 2021; Sporrij et al., 2023; Wichmann et al., 2022; Yamagata et al., 2021), whereas the more subtle drop in H4ac is consistent with known redundancies between NuA4/TIP60 and other HATs of the MYST family targeting histone H4 (Rea et al., 2007; Yokoyama et al., 2024).

**Figure 1.**
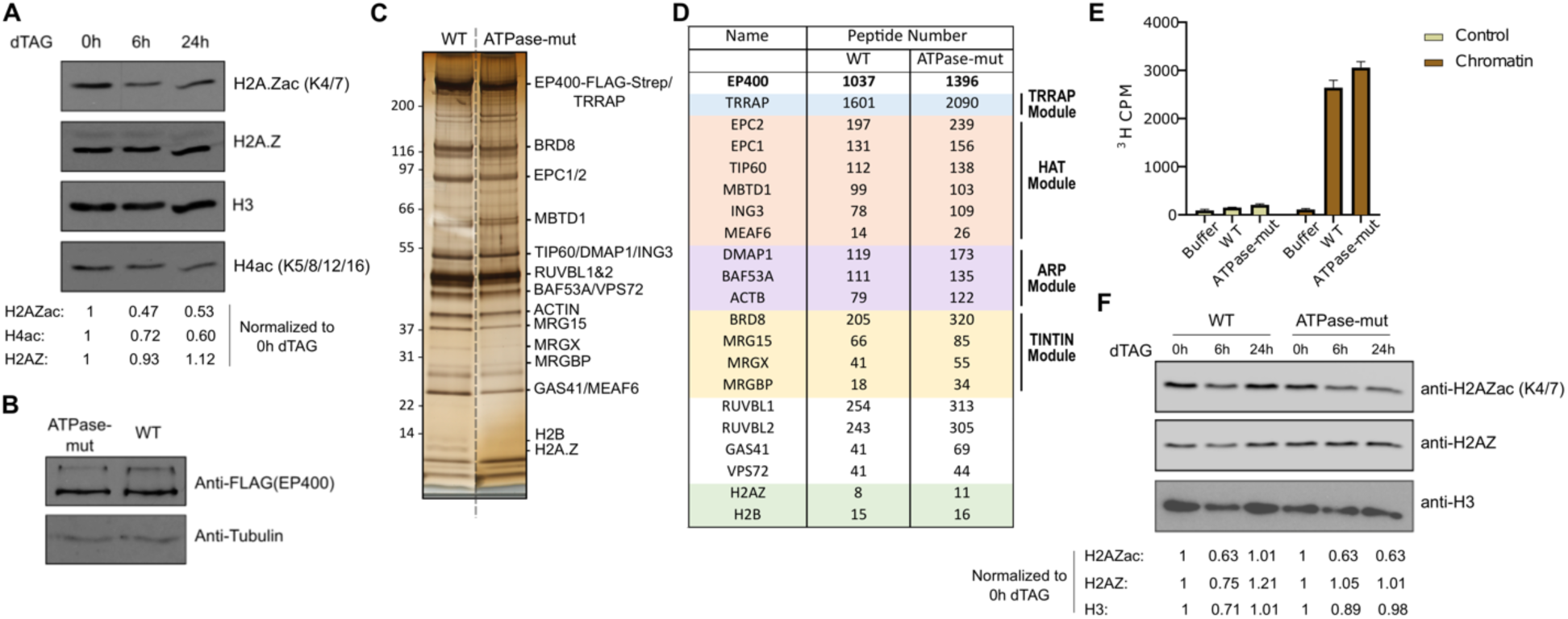
EP400 remodeling activity is crucial for maintaining acetylated H2A.Z on chromatin. **(A)** Western blots (WBs) on chromatin-enriched extracts (CEEs) derived from an EP400-dTAG cell line (clone 63) at 0h, 6h and 24h of dTAG treatment. **(B)** WB comparing protein expression of 3xFLAG-2xStrep-tagged WT and ATPase-mut EP400 in cell lines established through insertion of the 3xFLAG-2xStrep-tagged constructs into the AAVS1 locus of EP400-dTAG cells (clone 63). **(C)** Silver-stained gel of 3xFLAG-2xStrep-EP400 fractions (WT and ATPase-mut) obtained by tandem affinity purification. Biotin eluates (2^nd^ purification) were loaded on the gel. **(D)** Peptide counts of NuA4/TIP60 subunits and the H2A.Z-H2B dimer detected by mass spectrometry analysis of tandem affinity purified fractions. **(E)** HAT assay comparing acetyltransferase activities of WT and ATPase-mut fractions using native chromatin as a substrate and radio-labelled ^3^H-acetyl-CoA. The ^3^H counts per minute (CPM) are indicated on the y-axis. **(F)** WBs on CEEs derived from the cell lines in (B) at 0h, 6h and 24h of dTAG treatment.

To specifically interrogate the remodeling function of EP400, unconfounded by effects of its scaffolding role, we ectopically expressed 3x-FLAG-2xStrep-tagged wild type (WT) or K1085L mutant EP400 lacking ATPase activity (Xu et al., 2010) (ATPase-mut henceforth) from the *AAVS1* locus (Dalvai et al., 2015) in the dTAG system cells and selected clones with similar expression levels (Figure 1B). In these cells, dTAG-mediated EP400 knock-out is rescued by either WT or ATPase-mut EP400, which permits analysis of effects specific to loss of the remodeling function rather than complex disruption. To confirm that the ATPase-mut NuA4/TIP60 complex is otherwise intact, we performed tandem affinity purification of WT and ATPase-mut EP400. Mass spectrometry (MS) analysis validated full assembly of the complex, where we detected all the known subunits and modules in addition to the H2A.Z-H2B dimer in both samples to comparable levels (Figure 1C-D). We also confirmed retention of the HAT activity in the ATPase-mut complex by *in vitro* HAT assay (Figure 1E). As expected, no clear change was observed in total H2A.Z by Western blot analysis on CEEs in either WT- or ATPase-mut-complemented cells upon dTAG treatment (Figure 1F), similar to the non-complemented cells (Figures 1A and S1A-B). Interestingly, despite preservation of complex integrity and HAT activity in the ATPase-mut, we observed a clear drop in H2A.Zac in response to treatment (Figure 1F). These data suggest that the small fraction of chromatin-associated H2A.Z whose remodeling is dependent on EP400 consists of acetylated H2A.Z. One possible scenario is that Tip60 acetylates H2A.Z on chromatin following its incorporation by EP400 at NuA4/TIP60 target sites. Another possibility is that Tip60 pre-acetylates H2A.Z in the soluble fraction, resulting in H2A.Zac-H2B dimers which are then incorporated by EP400.

### Genome-wide EP400-dependent H2A.Z incorporation occurs at sites where H2A.Z is acetylated

To corroborate our conclusions from the bulk chromatin analysis, we analyzed genome-wide H2A.Z and H2A.Zac profiles by chromatin immunoprecipitation-sequencing (ChIP-seq) in WT-complemented (WT), non-complemented (dTAG) and ATPase-mut-complemented (ATPase-mut) cells treated with dTAG for 24h. The majority of H2A.Z peaks (∼74%) show no significant change in either dTAG or ATPase-mut conditions compared with the WT condition, with only ∼26% of the peaks showing a significant change in H2A.Z signal following remodeler loss (Figure 2A). By contrast, over half of H2A.Zac peaks (∼51%) significantly change following loss of remodeling activity (Figure 2B). These results are in line with our hypothesis of EP400 remodeling function being restricted to specific genomic regions, including sites of H2A.Z acetylation.

**Figure 2.**
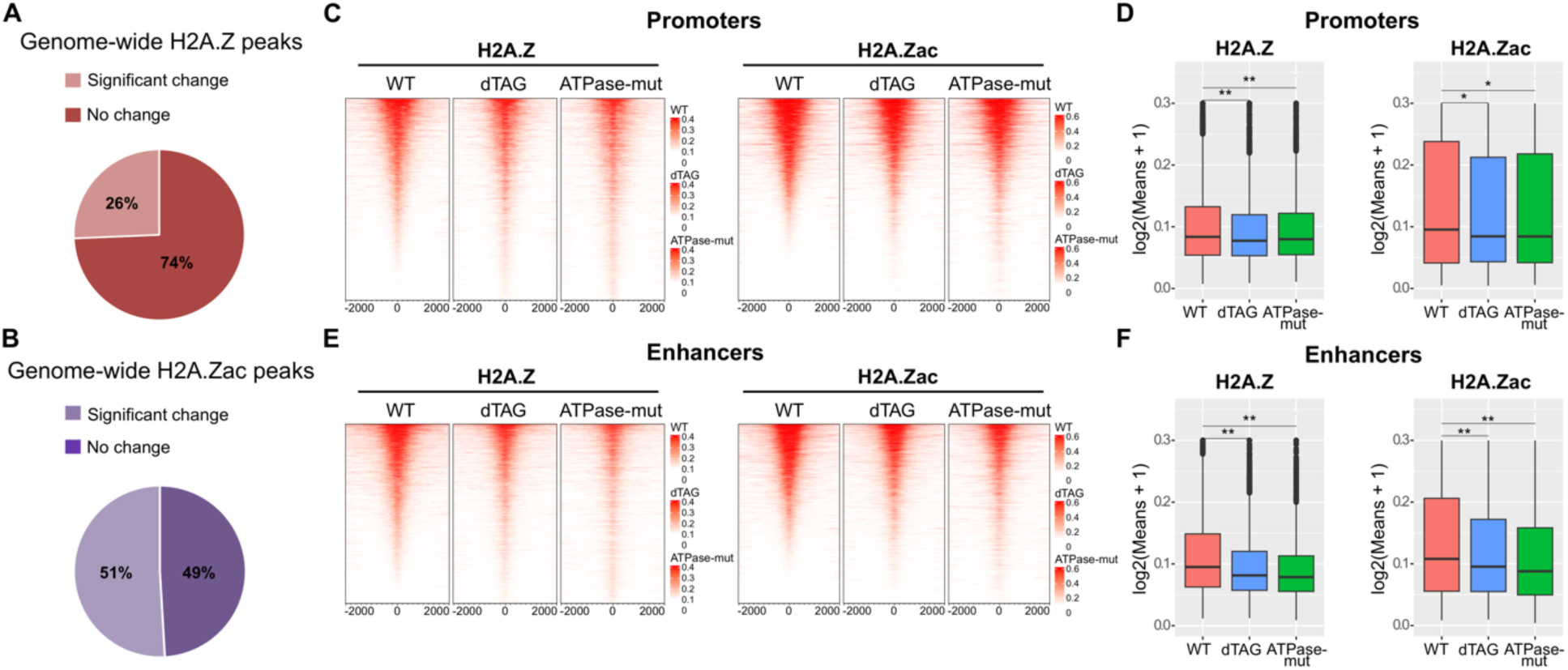
Genome-wide EP400-dependent H2A.Z incorporation occurs at sites where H2A.Z is acetylated. **(A-B)** Pie-charts representing the proportions of genome-wide **(A)** H2A.Z and **(B)** H2A.Zac peaks with significant signal change in dTAG or ATPase-mut relative to WT. **(C, E)** Left, Heatmap representation of H2A.Z peaks which significantly change in dTAG or ATPase-mut relative to WT, ranked in descending order based on signal in the WT condition at **(C)** promoters and **(E)** enhancers. Right, heatmap representation of H2A.Zac signal in WT, dTAG and ATPase-mut specifically for peaks which overlap the H2A.Z peaks depicted on the left panels. **(D)** Boxplot representation of the heatmaps in (C). **(F)** Boxplot representation of the heatmaps in (E). These ChIP-seq data are merged from two biological replicates (n=2). The read per million (RPM) signals were plotted on the heatmaps. Wilcoxon rank-sum test was performed on the boxplots (\**P* < 0.05, \*\**P* < 0.01).

To further investigate the change, we overlapped the significantly changing H2A.Z peaks with annotated promoters, enhancers, and other genomic regions. All regions show a significant mean decrease in H2A.Z signal (Figures 2C-F and S2A-F), consistent with loss of remodeling activity. Importantly, the significantly changing H2A.Z peaks largely overlap with H2A.Zac peaks (Figures 2C, 2E, S2A, S2C, and S2E), with the overlapping H2A.Zac signal also significantly decreasing at *cis*-regulatory elements (Figure 2C-F) and other genomic targets (Figure S2A-F). Taken together, these data strongly support an intimate link between H2A.Z incorporation by EP400 and its acetylation by Tip60 as a mechanism of chromatin modification and gene regulation.

### EP400 directly exchanges pre-acetylated H2A.Z at gene promoters

We next sought to confirm that sites where H2A.Z and H2A.Zac signals decrease in response to remodeler loss of function were direct targets of NuA4/TIP60, particularly at *cis*-regulatory elements. To this end, we mapped the genome-wide binding sites of EP400 and two other NuA4/TIP60 subunits, EPC1 and DMAP1, through ChIP-seq in K562 cell lines expressing endogenously FLAG-tagged EP400, EPC1 or DMAP1. When the peaks of each subunit are overlapped with H2A.Z peaks which significantly change in dTAG and ATPase-mut at promoters and enhancers (identified in Figure 2C and E), we see strongest enrichment of the NuA4/TIP60 complex at sites of highest H2A.Z signal (Figures 3A and S3), the same sites which show the most pronounced decrease in dTAG and ATPase-mut H2A.Z signal compared with WT. The enrichment is particularly high at promoters (Figure 3A) but much weaker at enhancers (Figure S3), suggesting that gene promoters constitute the main direct targets of EP400 remodeling activity.

**Figure 3.**
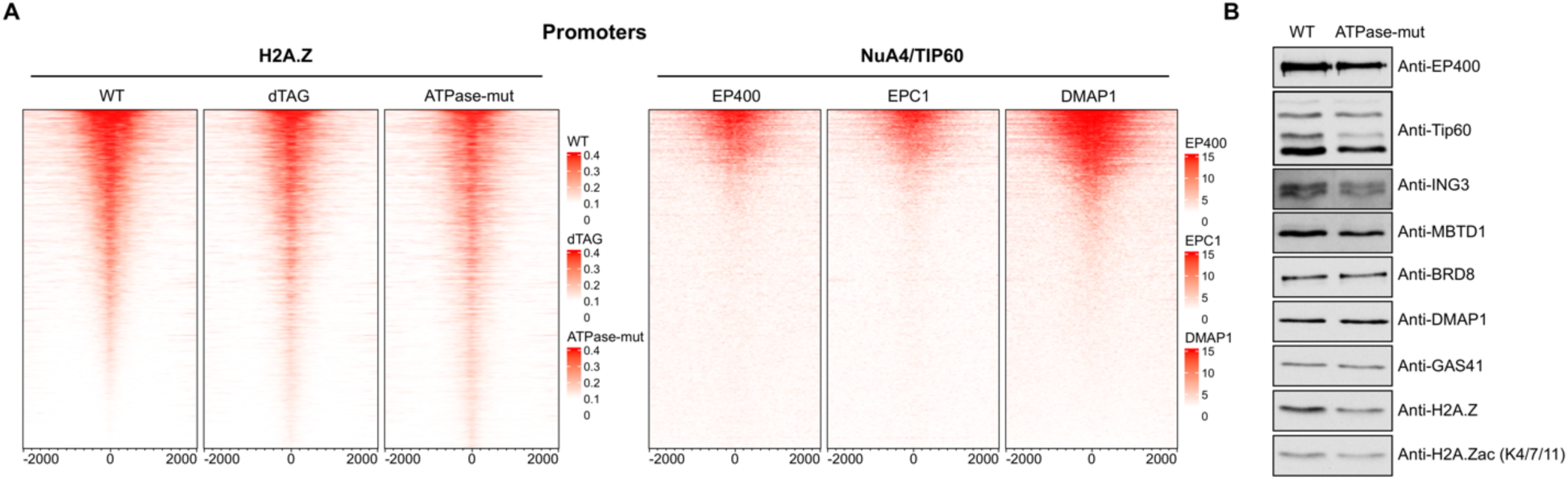
EP400 directly exchanges pre-acetylated H2A.Z at gene promoters. **(A)** Left, heatmap representation of promoter-specific significantly changing H2A.Z peaks (as previously shown on Figure 2C). Right, heatmap representation of EP400, EPC1 and DMAP1 peaks which overlap the H2A.Z peaks plotted on the left panel. **(B)** WB analysis performed on NuA4/TIP60 tandem affinity purified fractions from figure 1, detecting the indicated subunits, H2A.Z, and H2A.Zac.

We then verified if H2A.Z could be pre-acetylated by Tip60 before it is exchanged on chromatin by EP400 by checking the presence of H2A.Zac in a highly pure native NuA4/TIP60 fraction obtained by tandem affinity purification of EP400 from K562 soluble nuclear extract, i.e. excluding chromatin (see Figure 1C-D and Methods). Western blot analysis confirmed that H2A.Z within the H2A.Z-H2B dimer which is stably associated with the NuA4/TIP60 complex is acetylated (Figure 3B). Altogether, these data are in favor of a mechanism whereby EP400 directly deposits pre-acetylated H2A.Zac-H2B dimers at promoters of target genes to quickly activate or upregulate their transcription.

### NuA4/TIP60 positively regulates transcription of target genes through deposition of pre- acetylated H2A.Z at their promoters

To determine the functional relevance of our findings, we aimed to correlate chromatin alterations with transcriptional perturbations resulting from EP400 loss of function. First, we performed mRNA sequencing (RNA-seq) in the ATPase-mut-complemented and WT-complemented cell lines after 24h of dTAG treatment (Figure 4A-C). Differential gene expression analysis yielded 357 significantly downregulated genes and 808 significantly upregulated genes in the ATPase-mut vs the WT condition (Figure 4B-C). The top enriched biological pathways for the downregulated gene list include cell proliferation, and several cell fate and developmental processes including hematopoiesis and neurodevelopment (Figure 4D), consistent with reports on NuA4/TIP60-mediated H2A.Z remodeling or acetylation (Bellucci et al., 2013; Couture et al., 2012; Dalvai et al., 2013; Frob et al., 2019; Gevry et al., 2007; Janas et al., 2022; Jostes et al., 2024; Numata et al., 2020; Sporrij et al., 2023; Wichmann et al., 2022; Yang et al., 2024). By contrast, upregulated genes are enriched in processes which are not known targets of H2A.Z-specific NuA4/TIP60 function (Figure S4). These results suggest that, despite their lower number, downregulated genes in the EP400 loss of function mutant cells are the direct targets of EP400 remodeling activity, with gene upregulation being an indirect consequence of transcriptional dysregulation.

**Figure 4.**
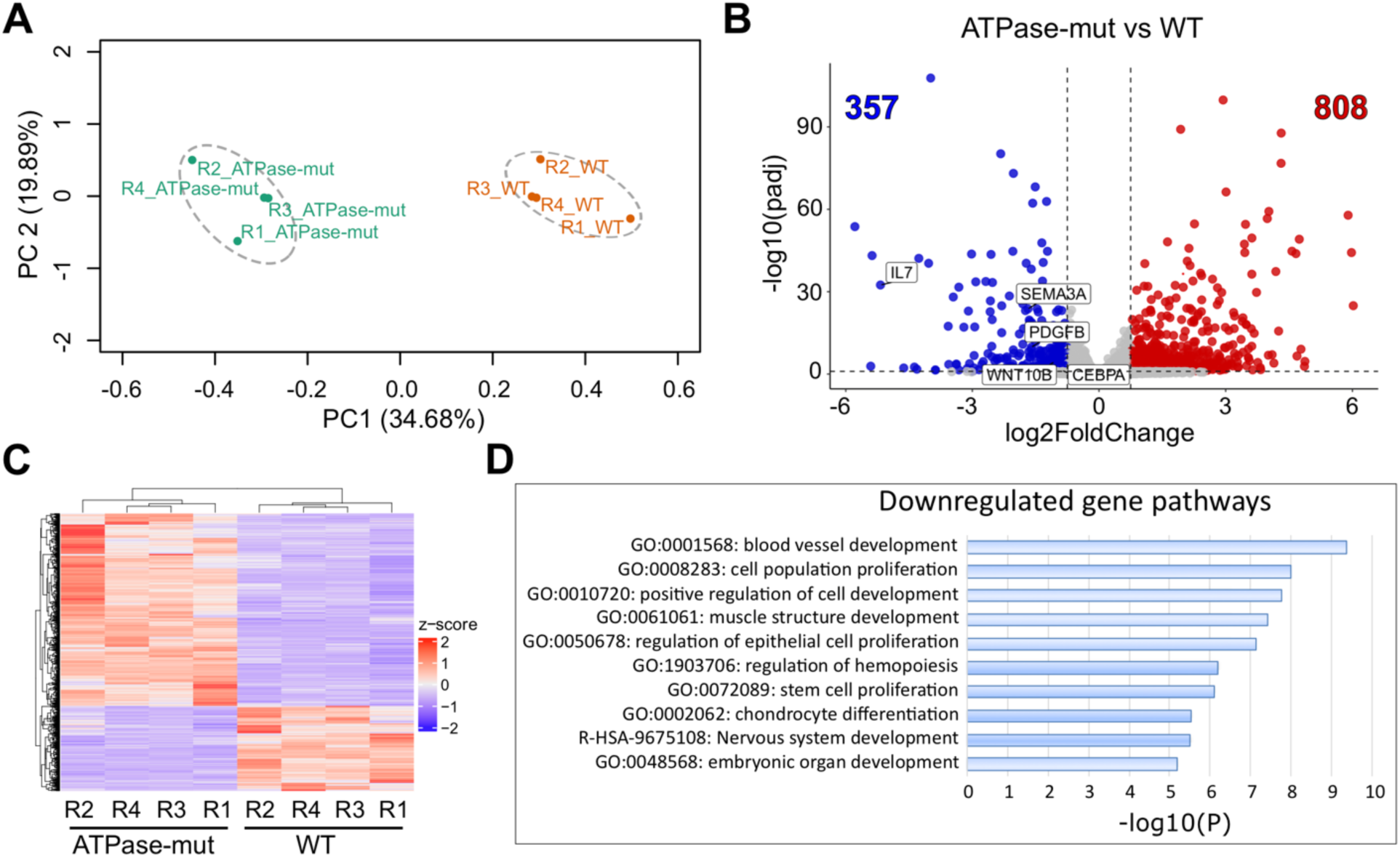
Transcriptional changes resulting from loss of EP400 ATPase activity. **(A)** PCA plot for WT-complemented and ATPase-mut-complemented RNA-seq samples. **(B)** Volcano plot representation of differential gene expression, with downregulated genes in blue (log2(FC) < - 0.75, padj < 0.1), upregulated genes in red (log2(FC) > 0.75, padj < 0.1), and unchanged genes in gray. Some of the downregulated genes are labelled on the graph and discussed in the context of specific pathways. Analysis was performed on four biological replicates (n=4) per condition. **(C)** Hierarchically clustered Z-score heatmap representing normalized transcript counts for ATPase- mut vs WT differentially expressed genes. **(D)** Pathway enrichment of downregulated genes identified in (B). Metascape (Zhou et al., 2019) was used to generate pathway enrichment according to the GO Biological Processes, Reactome Pathway, and KEGG Pathway databases. The 10 pathways plotted are among the top 20.

Next, we examined the change in H2A.Z and overlapping H2A.Zac signals in the dTAG and ATPase-mut conditions relative to the WT control specifically at transcription start site (TSS)- proximal regions of either downregulated or upregulated genes. Both H2A.Z and H2A.Zac significantly drop in both dTAG and ATPase-mut cells at downregulated gene promoters (Figures 5A-B, 5E-F and S5A-C), whereas no significant drop in either is seen at upregulated genes (Figure 5C-D). This is in agreement with the role of NuA4/TIP60 as a transcriptional co-activator and argues that EP400-mediated H2A.Z exchange at gene promoters can only be activating, and cannot be repressive, due to tight link to acetylation. Together, our results provide compelling evidence for positive transcriptional regulation of NuA4/TIP60 target genes through deposition of pre-acetylated H2A.Z by EP400 at their promoters.

**Figure 5.**
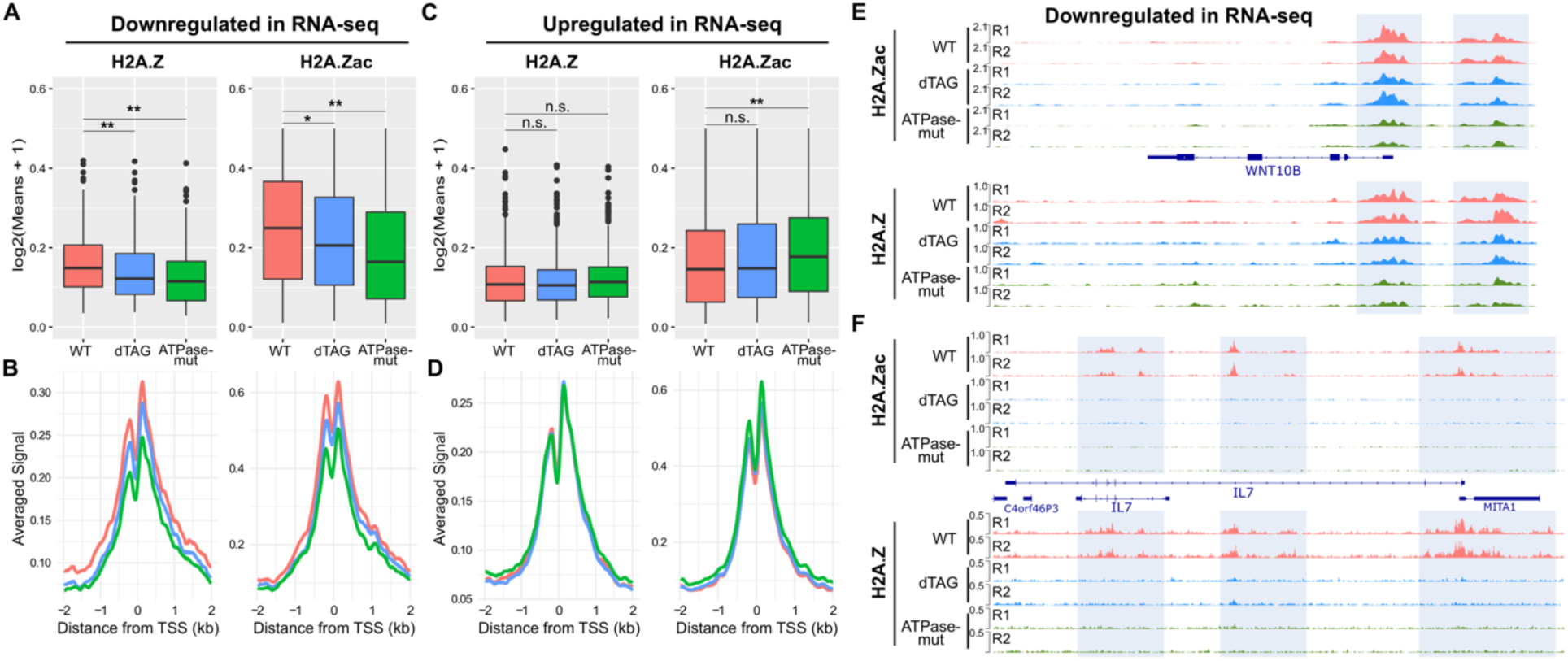
NuA4/TIP60 positively regulates transcription of target genes through deposition of pre-acetylated H2A.Z at their promoters. **(A, C)** Boxplot representation of H2A.Z peaks and the H2A.Zac peaks overlapping them at TSS-proximal regions (±2kb) of **(A)** genes identified as downregulated and **(C)** genes identified as upregulated in ATPase-mut vs WT RNA-seq (see Figure 4B). **(B, D)** Metaplots of the same peak signals represented in (A) and (C), showing averaged signal distribution around the TSS. **(E-F)** Integrative Genomics Viewer (IGV) snapshots of RPM-normalized profiles of H2A.Z and H2A.Zac at two genes, **(E)** WNT10B and **(F)** IL7, identified as downregulated in ATPase-mut vs WT RNA-seq. Wilcoxon rank-sum test was performed on the boxplots (\**P* < 0.05, \*\**P* < 0.01, n.s. = non-significant).

### NuA4/TIP60 HAT module is required for H2A.Z remodeling and transcriptional regulation

To gain further mechanistic insights into the remodeling function of the NuA4/TIP60 complex, we introduced mutations within the helicase-SANT-associated (HSA) domain of EP400, specifically in a region that we recently discovered to be important for interaction with a previously undescribed second copy of the subunit BAF53A. Component of the ARP modules of both NuA4/TIP60 and SRCAP, the first copy is important for nucleosome engagement by the remodeling complexes (Yang et al., 2024). We reasoned that the novel second copy may also play a role in modulating EP400 ATPase activity, and we decided to probe its function within the complex. First, two K562 cell lines were established through *AAVS1* expression of either a four residue-mutant 3xFLAG- 2xStrep-EP400 harboring the mutations I869D/A870D/A874D/I877D in the HSA domain (HSA- 4mut henceforth) or a six residue-mutant version containing the mutations I869D/A870D/A874D/I877D/F880D/W881D (HSA-6mut henceforth) (Yang et al., 2024). Next, to validate effects of the mutations on BAF53A copy number in the complex, we tandem affinity purified the HSA-4mut and HSA-6mut in parallel to WT EP400 and proceeded with MS analysis. To our surprise, the results show a drastic loss of the HAT module, composed of the subunits TIP60, EPC1/2, ING3, MEAF6 and MBTD1, in both HSA mutants (Figures 6A-C). We think that the mutations distorted the structure of the EP400 HSA domain, known to be a pillar for ARP module arrangement, which in turn is crucial for anchoring the HAT module to the rest of the complex. Another unexpected finding in our MS results is the lower detection of H2A.Z-H2B dimers in both HSA-mut fractions, despite the presence of VPS72, the chaperone responsible for binding the dimer to the complex (Yang et al., 2024), being unaffected by the mutation (Figure 6B). This suggests that pre-acetylation of H2A.Z by the HAT module is important for proper H2A.Z-H2B dimer association with the complex before remodeling. To further test this hypothesis and investigate the role of the HAT module and pre-acetylation in H2A.Z exchange, we inserted the HSA-6mut construct into the *AAVS1* locus of EP400-dTAG cells, and a clone with similar expression to the WT-complemented cell line was selected for downstream analysis (Figure 6D).

**Figure 6.**
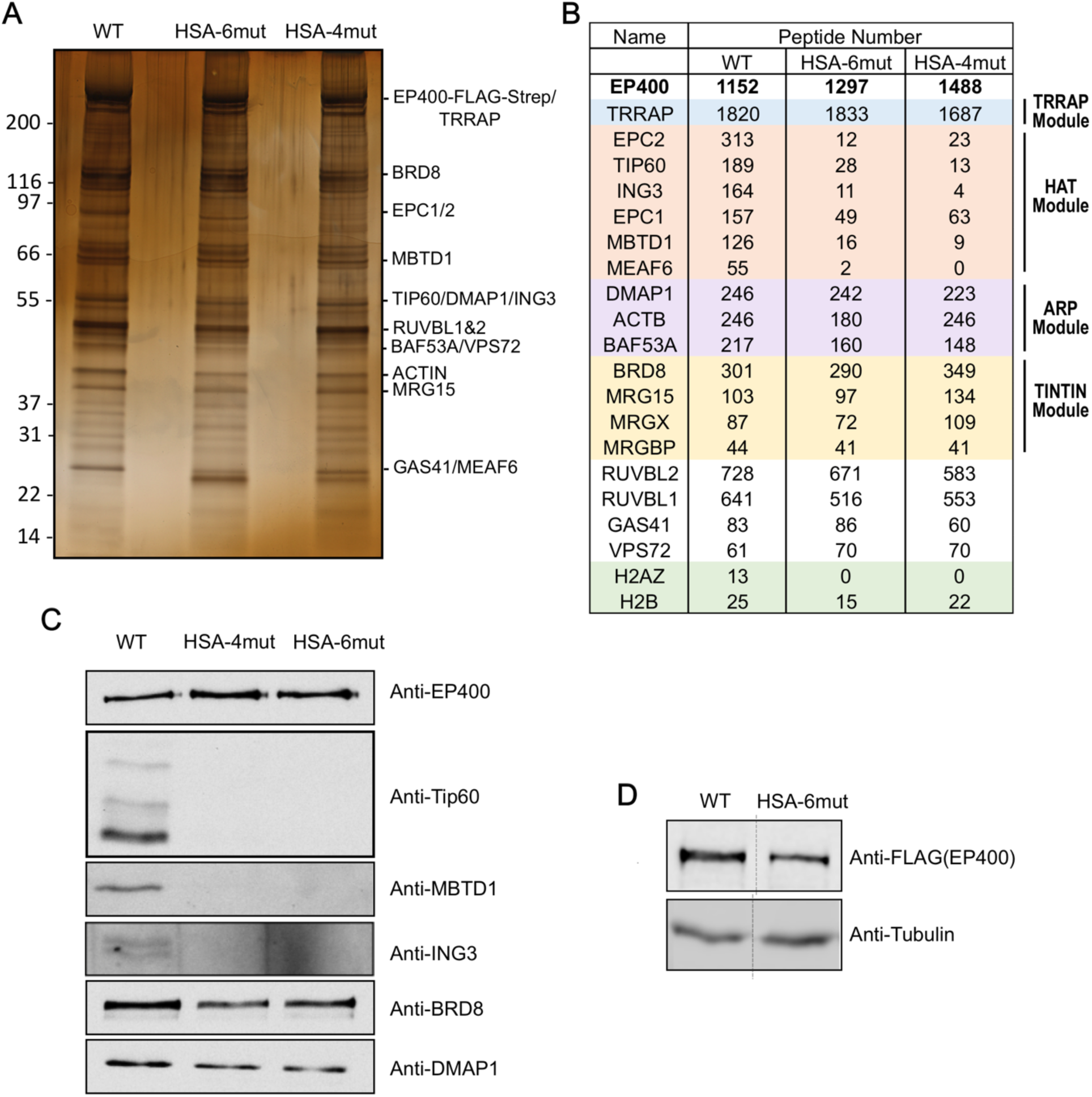
A model system for investigating the role of HAT activity in H2A.Z exchange by NuA4/TIP60. **(A)** Silver-stained gel of purified 3xFLAG-2xStrep-EP400 fractions (WT, HSA-6mut and HSA-4mut). FLAG eluates (1^st^ purification) were loaded on the gel. **(B)** Peptide counts of NuA4/TIP60 subunits and H2A.Z-H2B dimers detected by mass spectrometry analysis of tandem affinity purified fractions (biotin eluates/2^nd^ purification). **(C)** WB on the same purified fractions with the indicated antibodies for NuA4/TIP60 subunits. **(D)** WB comparing protein expression of 3xFLAG-2xStrep-tagged WT and HSA-6mut EP400 in cell lines established through insertion of the 3xFLAG-2xStrep-tagged constructs into the AAVS1 locus of EP400-dTAG cells (clone 63).

We then performed ChIP-seq of H2A.Zac and H2A.Z in HSA-6mut-complemented and WT-complemented dTAG cell lines following 24h of treatment. As expected from HAT module loss, H2A.Zac peaks which significantly change in the HSA-6mut relative to the WT mostly show reduced signal genome-wide (Figure S6A-B), including promoters and enhancers (Figure 7A-D). Interestingly, the overlapping H2A.Z peaks also show a significant drop in signal (Figures 7A-D and S6A-B), arguing that loss of H2A.Zac on chromatin in the absence of the HAT module is mainly due to lack of EP400-mediated H2A.Zac-H2B remodeling rather than loss of acetylation of pre- deposited H2A.Z on chromatin.

**Figure 7.**
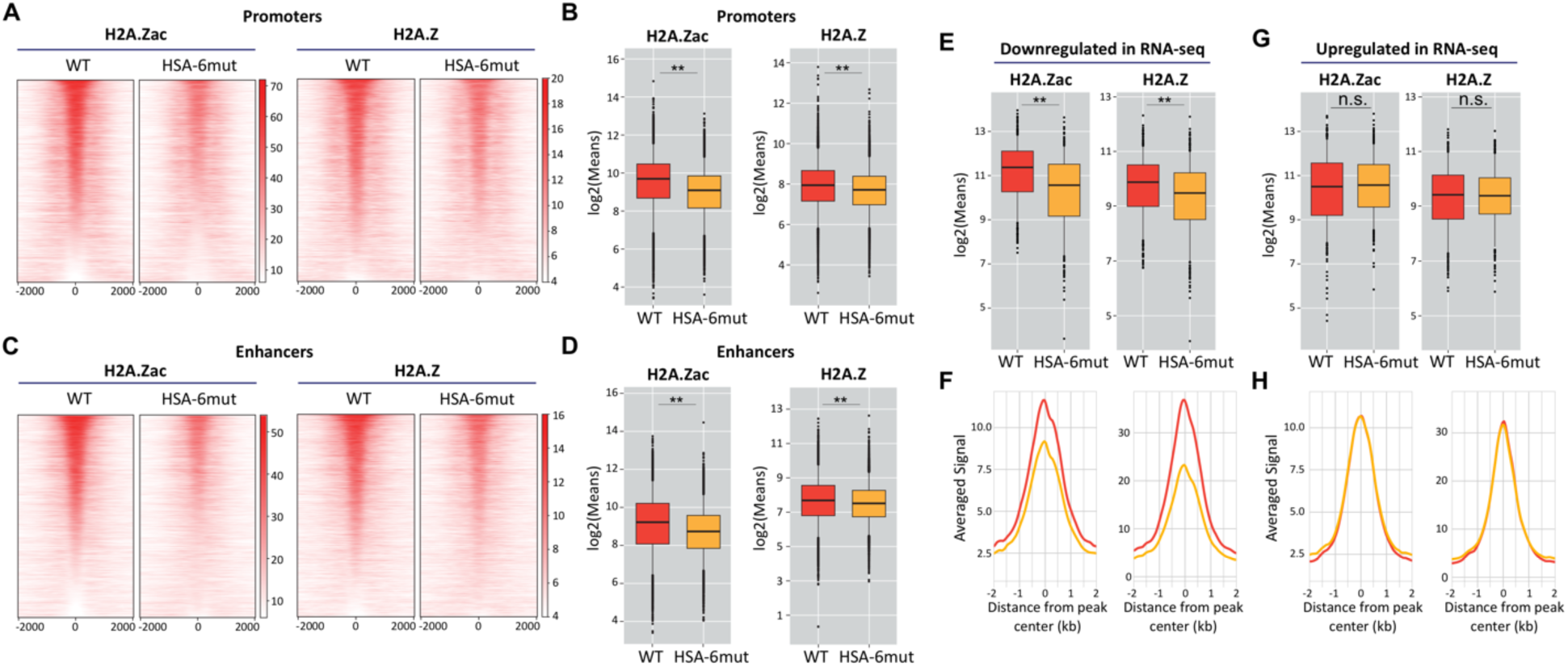
NuA4/TIP60 HAT module is required for H2A.Z remodeling and transcriptional regulation. **(A, C)** Left, Heatmap representation of H2A.Zac peaks which significantly change in HSA-6mut relative to WT, ranked in descending order based on signal in the WT condition at **(A)** promoters and **(C)** enhancers. Right, heatmap representation of H2A.Z signal in WT and HSA-6mut specifically for peaks which overlap the H2A.Zac peaks depicted on the left panels. **(B)** Boxplot representation of the heatmaps in (A). **(D)** Boxplot representation of the heatmaps in (C). **(E, G)** Boxplot representation of H2A.Zac peaks and the H2A.Z peaks overlapping them at TSS-proximal regions (±2kb) of **(E)** genes identified as downregulated and **(G)** genes identified as upregulated in HSA-6mut vs WT RNA-seq (based on Figure S7). **(F, H)** Metaplots of the same peaks represented in (E) and (G), showing averaged signal across all peaks. These ChIP-seq data are merged from two biological replicates (n=2). The read per million (RPM) signals were plotted on the heatmaps. Wilcoxon rank-sum test was performed on the boxplots (\*\**P* < 0.01, n.s. = non-significant).

We further correlated chromatin changes with transcription by first performing RNA-seq in the HSA-6mut compared with WT cells (Figure S7A-C). Both H2A.Zac peaks and the overlapping H2A.Z signal found in TSS-proximal regions of downregulated genes exhibit a pronounced decrease in the HSA-mut relative to WT cells (Figure 7E-F), further confirming that the HAT module is crucial for H2A.Z exchange by the NuA4/TIP60 complex at promoters of its target genes. Upregulated genes on the other hand show no significant change in either H2A.Zac or H2A.Z (Figure 7G-H). Overall, our data argue for a model in which H2A.Z-H2B dimers are first targeted by Tip60 which acetylates H2A.Z, in cis within the NuA4/TIP60 complex or in trans allowing for H2A.Zac-H2B association. ATP-dependent remodeling by EP400 then allows H2A.Zac incorporation in chromatin, particularly at gene promoters to positively regulate transcription.

## DISCUSSION

The roles of the NuA4/TIP60 complex in either H2A.Z remodeling (Bellucci et al., 2013; Bowman et al., 2011; Couture et al., 2012; Frob et al., 2019; Gevry et al., 2007; Jostes et al., 2024; Pradhan et al., 2016; Sudarshan et al., 2022; Sun et al., 2023; Yang et al., 2024) or H2A.Z acetylation (Dalvai et al., 2013; Giaimo et al., 2018; Ito et al., 2018; Janas et al., 2022; Numata et al., 2020; Procida et al., 2021; Sporrij et al., 2023; Wichmann et al., 2022; Yamagata et al., 2021) have been repeatedly described in the context of transcriptional regulation. However, the two enzymatic functions targeting the same histone variant have thus far been studied separately, with the coordination between them remaining unexplored despite large overlap of key physiological and pathological processes regulated by either function. With our EP400 rapid degradation system, complemented by WT or mutant versions of the protein, we managed to model specific loss of either ATPase activity or HAT activity of NuA4/TIP60 and untangle the enzymatic interdependence which exists within the complex through functional genomic analyses.

We uncover a critical role for the remodeling activity of EP400 in maintaining acetylated H2A.Z on chromatin, hinting at the importance of this remodeler in specifically depositing H2A.Zac at target genomic sites. Indeed, we find that about a quarter of H2A.Z-enriched genomic sites in K562 cells depend on EP400 action to maintain H2A.Z binding, and that those sites coincide with regions where H2A.Z is acetylated and where H2A.Zac signal also drops upon ATPase loss of function. Further, our data strongly support a role for EP400 in incorporating pre- acetylated H2A.Z at target gene promoters, positively regulating transcriptional activity, particularly at genes involved in pathways which have already been linked to either Tip60- mediated H2A.Z acetylation or EP400-mediated H2A.Z remodeling, including regulation of cell proliferation (Bellucci et al., 2013; Dalvai et al., 2013; Gevry et al., 2007; Jostes et al., 2024; Wichmann et al., 2022; Yang et al., 2024), cell fate and development (Couture et al., 2012; Frob et al., 2019; Janas et al., 2022; Numata et al., 2020; Sporrij et al., 2023), and cancer (Ito et al., 2018; Jostes et al., 2024; Sun et al., 2023; Yamagata et al., 2021). In turn, pre-acetylation seems to be important for H2A.Z-H2B exchange by EP400, as we find that upon loss of the NuA4/TIP60 HAT module, genomic sites with decreased H2A.Zac signal also show a significant decrease in total H2A.Z, which means that remodeling is impaired in the absence of HAT activity. Although it is possible that loss of NuA4/TIP60-mediated H2A and H4 acetylation on chromatin in the absence of the HAT module affects H2A.Z remodeling efficiency to some extent, as has been described in lower eukaryotes (Altaf et al., 2010), we find evidence that pre-acetylation of H2A.Z may be important for full association of the H2A.Z-H2B dimer with the complex, arguing that pre- acetylation is a key step preceding incorporation of the variant into chromatin by EP400.

In the ATPase-mut condition, we see a slight increase in H2A.Zac at upregulated genes while H2A.Z remains unchanged (Figures 5C-D). This suggests that the HAT-competent ATPase mutant NuA4/TIP60 complex may acetylate pre-deposited H2A.Z at promoters of upregulated genes in an attempt to compensate for effects at downregulated genes. Incorporation of H2A.Z at such sites may be dependent on SRCAP. Indeed, NuA4/TIP60 is capable of acetylating nucleosomal H2A.Z *in vitro* (Figure S8). Therefore, incorporation of pre-acetylated H2A.Z by EP400 as a regulatory mechanism at some genomic loci does not exclude the possibility of acetylation of chromatin-bound H2A.Z by NuA4/TIP60 at other sites or under certain circumstances.

The interdependence between remodeling and acetyltransferase activities within the NuA4/TIP60 complex distinguishes EP400 from SRCAP as a remodeler of H2A.Z, where loading of pre-acetylated H2A.Z on chromatin is presumably a mechanism that is specific to EP400. However, as discussed above, cooperation between SRCAP and the NuA4/TIP60 HAT activity at certain genomic regions remains a possibility. This is supported by reports on SRCAP acting as a co-activator through H2A.Z deposition at target gene promoters (Cuadrado et al., 2010; Ding et al., 2024; Greenberg et al., 2019; Pugacheva et al., 2024; Slupianek et al., 2010; Sun et al., 2022; Wong et al., 2007; Xu et al., 2021; Yang et al., 2012; Ye et al., 2017; Zhang et al., 2024; Zhao et al., 2019), suggesting that the deposited H2A.Z is acetylated, which leads to positive transcriptional regulation. Further work comparing the genome-wide functions of the two complexes is necessary to establish the cooperation, redundancy, and/or distinct functions that might exist.

Although we have only explored the coordination between enzymatic activities of the NuA4/TIP60 complex in K562 cell lines, genes that are dependent on H2A.Zac loading by EP400 for transcriptional activity are involved in pathways that are reminiscent of known H2A.Z-targeted functions of NuA4/TIP60 as reported in studies in several model systems. Consequently, we are confident that the conceptual findings in our study on coordination of function within NuA4/TIP60 are applicable to other models. Nevertheless, it is important to study specific pathways in more physiologically relevant contexts. For examples, the hematopoiesis genes CEBPA and IL7 (Figures 5F and S5A), and the neurodevelopmental gene SEMA3A (Figure S5C) are all downregulated following a decrease in H2A.Z(ac) incorporation in the ATPase-mut condition. Moreover, the embryonic development gene WNT10B (Figures 5E) and the cell proliferation gene PDGFB (Figure S5B), which are both linked to cancer, are also downregulated and show reduced H2A.Z(ac) deposition in the remodeling mutant. As previously described, all of these pathways have been linked to either H2A.Z acetylation or exchange by NuA4/TIP60. Further investigation is required to establish the role of concomitant H2A.Z acetylation and exchange in each of these pathways in appropriate models.

The concerted action of HAT and ATPase activities within NuA4/TIP60 reported herein is particularly relevant to diseases such as cancer (Ito et al., 2018; Jostes et al., 2024; Sun et al., 2023; Yamagata et al., 2021). Selective small molecule inhibitors have been described for Tip60 and used in pre-clinical trials in various cancers (Zohourian et al., 2024). Combinatorial targeting of both the Tip60 and EP400 enzymes, especially in cancers with altered H2A.Z(ac) profiles, presents a viable therapeutic strategy which may further improve outcome.

## ACKNOWLEDGEMENTS

This work was supported by grants from the Canadian Institutes of Health Research (CIHR) to J.C. (FDN-143314, PJT-183708). A.M. held studentships from the Natural Sciences and Engineering Research Council of Canada and the Fonds de Recherche du Québec-Santé. J.C. held the Canada Research Chair in Chromatin Biology and Molecular Epigenetics.

## AUTHOR CONTRIBUTIONS

A.M., C.L., J.H., A.L. and J.P.L. performed the experiments. J.P.L., A.D., S.M.I.H. and J.C. supervised and secured funding. C.C., S.B., C.J.B., E.T., A.B. and S.M.I.H. analyzed the genomic data. A.M. and J.C. wrote the manuscript with the help of co-authors.

## METHODS

### Data availability

ChIP-seq and RNA-seq alignment files will be accessible on the Gene Expression Omnibus (GEO) database and raw mass spectrometry data will be accessible on the MassIVE database at the time of publication.

### Cell line construction and culture

The K562 EP400 acute knock-out (EP400-dTAG) cell lines and the WT-complemented dTAG cell line used in this study were produced in previous work (Yang et al., 2024). The ATPase-mut- and HSA-6mut-complemented dTAG cell lines were established by inserting 3xFLAG-2xStrep-tagged constructs into the *AAVS1* safe harbor locus of the EP400 dTAG clone 63 as previously described (Dalvai et al., 2015), and clones with similar expression to the WT-complemented cell line were selected and used in experiments (Figures 1B and 6C). Clonal expression was checked by Western blots on whole cell extracts using the following antibodies at the indicated dilutions: anti-FLAG Sigma A8592 (1:10000) and anti-Tubulin (alpha) Millipore CP06 (1:1000). K562 cell lines with *AAVS1* expression of 3xFLAG-2xSterp-tagged WT, HSA-6mut or HSA-4mut EP400 which were used in tandem affinity purification (Figures 6A-B) were generated in previous work (Yang et al., 2024). K562 cell lines with endogenously tagged EPC1 or EP400 (3xFLAG-2xStrep) were engineered by (Dalvai et al., 2015), and the DMAP1 endogenously tagged cell line was similarly constructed, as described in (Agudelo et al., 2017). All cell lines were cultured at 37°C with 5% CO_2_ in RPMI 1640 medium supplemented with 10% newborn calf serum and 1% GlutaMax.

### Tandem affinity purification

Tandem affinity purification of WT or mutant 3xFLAG-2xStrep-tagged EP400 was performed to obtain native complexes from nuclear extracts as previously described (Doyon & Cote, 2016). For the ATPase-mut vs WT purifications, biotin eluates (second purification) were used for the silver-stained gel, mass spectrometry analysis and HAT assay (Figures 1C-E). The biotin eluate of the WT complex was also used for examining H2A.Z acetylation within the complex by Western blot (Figure 3B), where the following antibodies were used: anti-H2A.Z Abcam ab4174 (1:1000) and anti-H2A.Zac (K4, K7, K11) Diagenode C15410173 (1:1000). For the HSA-6mut/HSA-4mut vs WT purifications, FLAG eluates (first purification) were used to produce the silver-stained gel, while biotin eluates were used for mass spectrometry analysis. Antibodies for different NuA4/TIP60 subunits used in immunoblots include: EP400 Abcam ab5201, Tip60/KAT5 Santa Cruz Biotechnology sc-5727, DMAP1 Thermo Fisher Scientific PA1-886, ING3 Abcam ab3713, BRD8 Bethyl Laboratories A300-219A, GAS41/YEATS4 Santa Cruz Biotechnology sc-393708, MBTD1 Abcam ab116361, FLAG Sigma A8592.

### Mass spectrometry

Mass spectrometry analysis was conducted at the Proteomics Platform of the Quebec Genomic Center using an Orbitrap Fusion mass spectrometer (Thermo Fisher Scientific) as detailed in (Devoucoux, Roques, et al., 2022). The raw data have been made public as indicated under “Data availability”.

### Chromatin-enriched extract (CEE) analysis

Non-complemented and WT- or ATPase-mut-complemented EP400-dTAG K562 cell lines were treated with 500 nM dTAG^V^-1 for 6h or 24h, or with DMSO for 24h as a control (0h condition). 4x10^6^ cells per condition were pelleted (1000xg, 3 min, 4°C) and washed twice with PBS, then lysed in 300 μL CEE Lysis Buffer (50 mM Tris pH 7.5, 100 mM NaCl, 0.5% NP-40, 1 mM EDTA, 1 mM DTT, 1 mM PMSF, 10 mM Sodium butyrate, 10 mM Nicotinamide, 10 mM Beta-glycerophosphate, 2 ug/mL Leupeptine, 2 ug/mL Pepstatine A, 5 ug/mL Aproptinine) and incubated on ice for 5 min. Lysates were pelleted (1000xg, 5 min, 4°C) and washed once with 100 μL CEE Digestion Buffer (50mM Tris pH 7.5, 300mM NaCl, 5 mM CaCl_2_, 1 mM DTT, 1 mM PMSF, 10 mM Sodium butyrate, 10 mM Nicotinamide, 10 mM Beta-glycerophosphate, 2 ug/mL Leupeptine, 2 ug/mL Pepstatine A, 5 ug/mL Aproptinin), then resuspended again in 100 μL CEE Digestion Buffer in addition to 4 U of micrococcal nuclease. Samples were incubated at 30°C in Eppendorf Thermomixer with 1200 rpm shaking for 40 min to solubilize chromatin, then centrifuged at 16000xg for 15 min at 4°C. The supernatants were collected as CEE and dosed with Bradford Protein Assay. 2 μg of each was loaded on 15% SDS-PAGE gels for Western blot analysis. The following antibodies were used for immunoblotting at the indicated dilutions: anti-H2A.Z Abcam ab4174 (1:2000), anti-H2A.Zac (K4, K7) Cell Signaling Technology 75336S (1:5000), anti-H3 Abcam ab1791 (1:20000), and anti-H4ac (K5, K8, K12, K16) Active Motif 39925 (1:5000).

### Histone acetyltransferase (HAT) assays

Gel fluorography and in-solution radioactivity count HAT assays were performed as detailed in (Barry et al., 2022). In the HAT assay comparing activities of the ATPase-mut and the WT complex fractions, the substrate used consists of short oligonucleosomes prepared as previously described (Utley et al., 1996). The substrates used in the gel-based HAT assay are recombinant mononucleosomes containing the indicated histones (H2A nucleosome EpiCypher 16-0006, H2A.Z.1 nucleosome EpiCypher 16-0014, and H2A.Z.2 nucleosome EpiCypher 16-0015).

### ChIP-seq

A detailed protocol for chromatin extraction, sonication, and immunoprecipitation can be found in (Yang et al., 2024). Briefly, two biological replicates of EP400-dTAG complemented and non-complemented cell lines were first treated with 500 nM dTAG^V^-1 for 24 h, following which chromatin was extracted and sheared into ∼200 bp fragments by sonication. The endogenously tagged EP400, EPC1 and DMAP1 cell lines were directly processed for ChIP with no treatment. For the dTAG cell lines, 100 ug of chromatin per replicate was immunoprecipitated with either 3 ug anti-H2A.Z (Abcam ab4174) or 0.2 ug anti-H2A.Zac (Diagenode C15410173). For the NuA4/TIP60 subunits ChIP-seq, 500 ug of chromatin per cell line was immunoprecipitated with either 3 ug anti-FLAG (Sigma F1804) or 3 ug IgG (rabbit, Millipore PP64). KAPA Hyper Prep kit (Roche, 07962363001) was used for library construction according to manufacturer’s protocol, and the samples were sequenced on Illumina NovaSeq 6000 with 100bp paired-end reads. HSA- 6mut vs WT data were analyzed and the associated heatmaps were plotted as previously described (Yang et al., 2024). Associated boxplots were generated with Python (v3.11.5) (Van Rossum, 2009), using Pandas (v2.0.3), Matplotlib (v3.7.2), Seaborn (0.12.2), SciPy (1.11.1) and NumPy (v1.24.3) libraries, averaging deepTools computeMatrix output values across peak coordinates. Statistical significance of the boxplots was evaluated with two-sided Mann-Whitney U test corrected by Benjamini Hochberg adjustment. For the rest of the data, raw reads were trimmed using fastp v0.21.0 (Chen et al., 2018). Trimmed reads were aligned on the human genome (hg38) using bwa mem v0.7.17 (Li, 2013) and samtools v1.13 (Li et al., 2009). Raw signal tracks and normalized tracks (in reads per million/RPM) were produced from mapped reads using deepTool’s bamCoverage tool v3.5.0 (Ramirez et al., 2014) and bedtools genomecov tool v2.27.1 (Quinlan & Hall, 2010), respectively. Tracks were converted to the bigwig format using bedGraphToBigWig v4 (Kent et al., 2010) and signal tracks were visualized with IGV (Robinson et al., 2011). The macs3 v3.0.0a7 (Feng et al., 2012) software was used to perform the peak calling relative to control (input for H2A.Z and H2A.Zac ChIP-seq, IgG for FLAG ChIP-seq) and regions were annotated using the ChIPseeker package v1.30.3 (Yu et al., 2015) in R v4.1.2 (R-Core-Team, 2018). The read count table was generated for counts within the peaks (summit 500 bp) and differential binding analysis was then performed using the DESeq2 v1.40.2 package (Love et al., 2014). Significantly changing peaks (padj < 0.2, fold change = 1.3) were divided into groups according to their genomic distribution for the generation of heatmaps using ComplexHeatmap v2.18.0 (Gu et al., 2016) and EnrichedHeatmap v1.32.0 (Gu et al., 2018) packages. The annotation of the enhancers was carried out according to the K562 enhancers known from EnhancerAtlas 2.0 (Gao & Qian, 2020) after conversion to hg38 genome with liftOver rtracklayer package function 1.62.0 (Lawrence et al., 2009). Boxplots and associated Wilcoxon tests were computed using means of bins per regions and were produced using the ggboxplot and stat_compare_means functions, respectively, with the default settings of the ggpubr v0.6.0 package (Kassambara, 2023). Metaplots were generated using the ggpubr v0.6.0 package by averaging the signal across all features for each bin of 100 bp. Promoter regions of significantly up- or downregulated genes (see “RNA-seq” below) were determined around TSSs (±2kb) and then overlapped with significantly changing peaks of H2A.Z and H2A.Zac to create boxplots and metaplots based on the signal of H2A.Z or H2A.Zac.

### RNA-seq

Cell line treatment, RNA extraction and library preparation were carried out as described (Yang et al., 2024). For the HSA-mut vs WT experiment, 3 biological replicates per condition were processed. Two experiments with biological duplicates each of ATPase-mut and WT samples were performed (for a total of 4 biological replicates per condition). HSA-mut vs WT analysis was performed as described (Yang et al., 2024). For the ATPase-mut vs WT analysis, reads were trimmed using fastp v0.23.2 (Chen et al., 2018). Quality check was performed on raw and trimmed data to ensure the quality of the reads using FastQC v0.11.9 (Andrews, 2010) and MultiQC v1.12 (Ewels et al., 2016). The quantification was performed with Kallisto v0.48.0 (Bray et al., 2016) against the human reference transcriptome (GRCh38/hg38 downloaded from Ensembl release 109). Batch effects of library preparation and sequencing were removed using a set of “in silico” empirical control genes and the RUVseq v1.34.0 package (Risso et al., 2014) considering k=1 factors of unwanted variation. The PCA was created by the plotPCA() function of the EDASeq v2.34.0 package (Risso et al., 2011) and the Volcano plot with the ggplot2 v3.4.4 package (Wickham, 2011). Differential expression analysis was performed using the DESeq2 v1.40.2 package (Love et al., 2014). To consider a differential gene expression statistically significant, thresholds of 0.1 and absolute log2(FC) = 0.75 were applied for the adjusted p-value and the minimum fold-change, respectively (as shown on the Volcano plot and heatmap). Heatmaps were generated using the ComplexHeatmap v2.18.0 package (Gu et al., 2016). All R analyses were done in R v4.3.1 (R-Core-Team, 2018). Pathway enrichment was performed using Metascape (Zhou et al., 2019).

---

**Figure S1.**
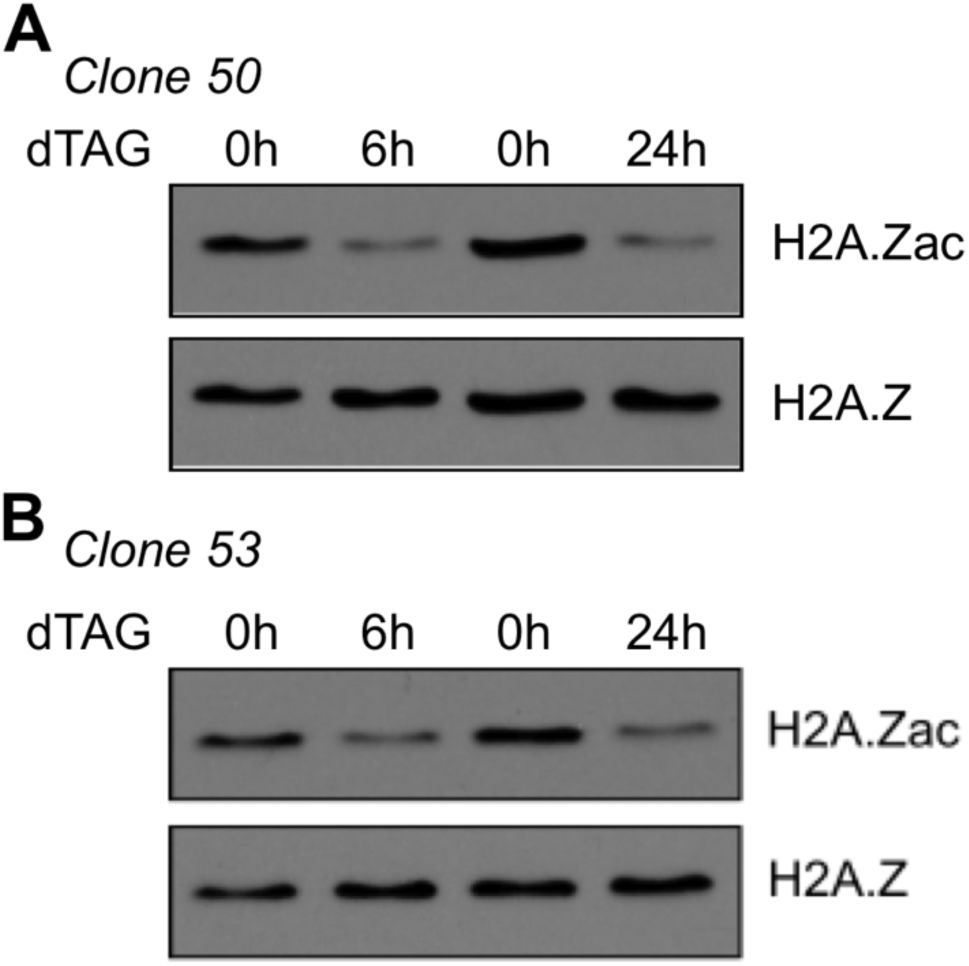
CEE analysis in EP400-dTAG cell lines. WB analysis of CEEs derived from two EP400-dTAG cell lines: **(A)** clone 50 and **(B)** clone 53, at 3 timepoints of dTAG treatment.

**Figure S2.**
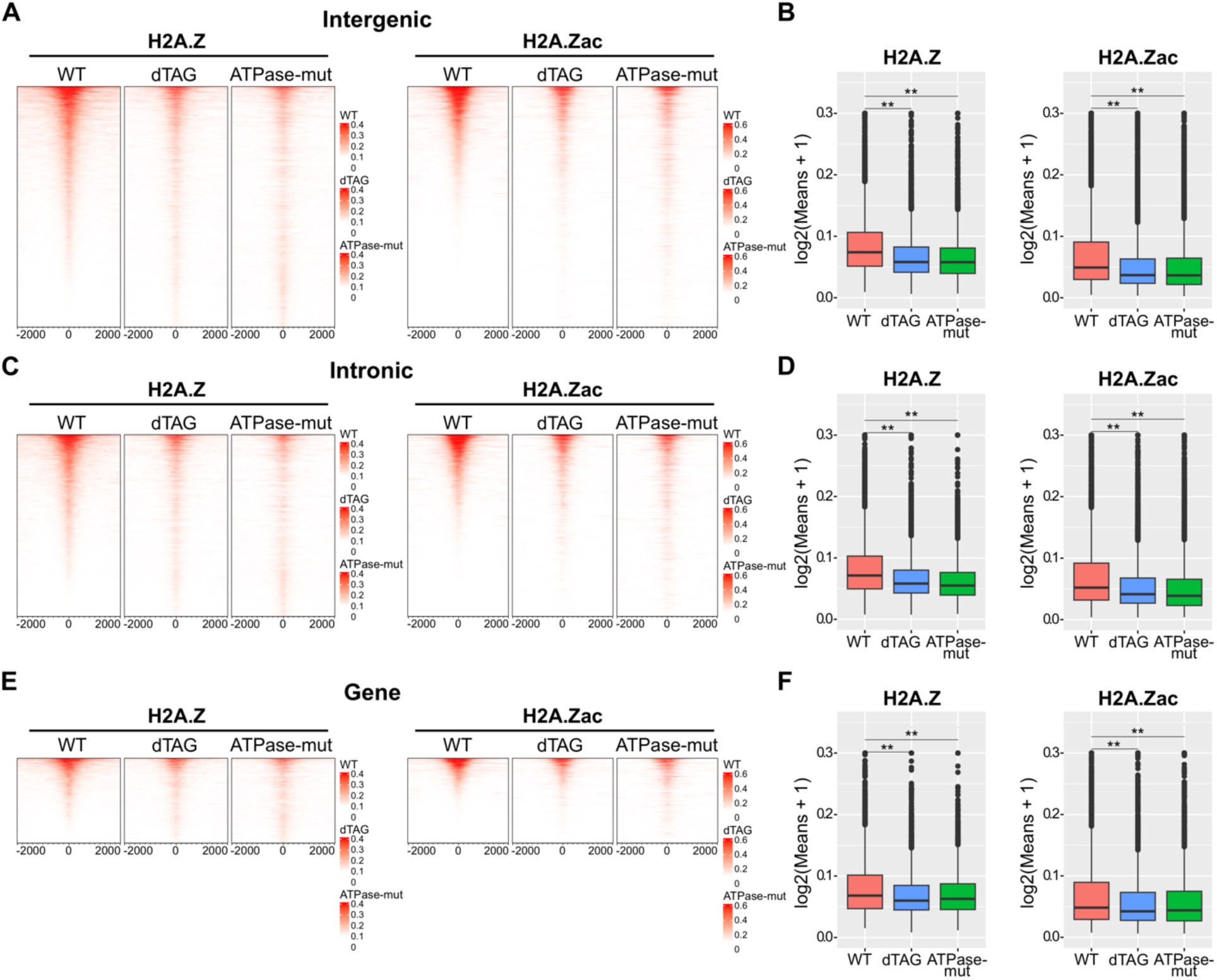
Genome-wide EP400-dependent H2A.Z incorporation occurs at sites where H2A.Z is acetylated. Related to Figure 2. **(A, C, E)** Left, Heatmap representation of H2A.Z peaks which significantly change in dTAG or ATPase-mut relative to WT, ranked in descending order based on signal in the WT condition at **(A)** intergenic regions, **(C)** intronic regions and **(E)** and genic regions that are either exons, 5’UTRs or 3’UTRs. Right, heatmap representation of H2A.Zac signal in WT, dTAG and ATPase-mut specifically for peaks which overlap the H2A.Z peaks depicted on the left panels. **(B, D, F)** Boxplot representations of the heatmaps in (A), (C) and (E), respectively. ChIP-seq data are merged from two biological replicates (n=2). The read per million (RPM) signals were plotted on the heatmaps. Wilcoxon rank-sum test was performed on the boxplots (\*\**P* < 0.01).

**Figure S3.**
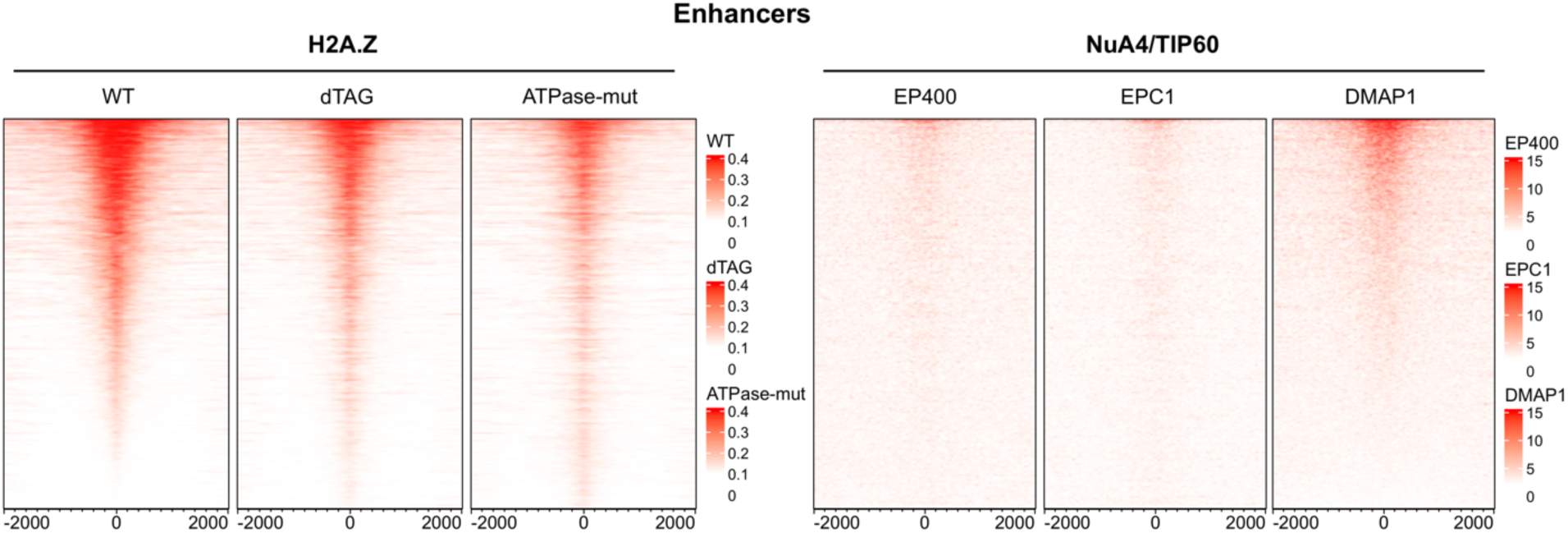
NuA4/TIP60 binding at enhancers that show significant H2A.Z(ac) change upon remodeler loss. Left, heatmap representation of enhancer-specific significantly changing H2A.Z peaks (as previously shown on Figure 2C). Right, heatmap representation of EP400, EPC1 and DMAP1 peaks which overlap the H2A.Z peaks plotted on the left panel.

**Figure S4.**
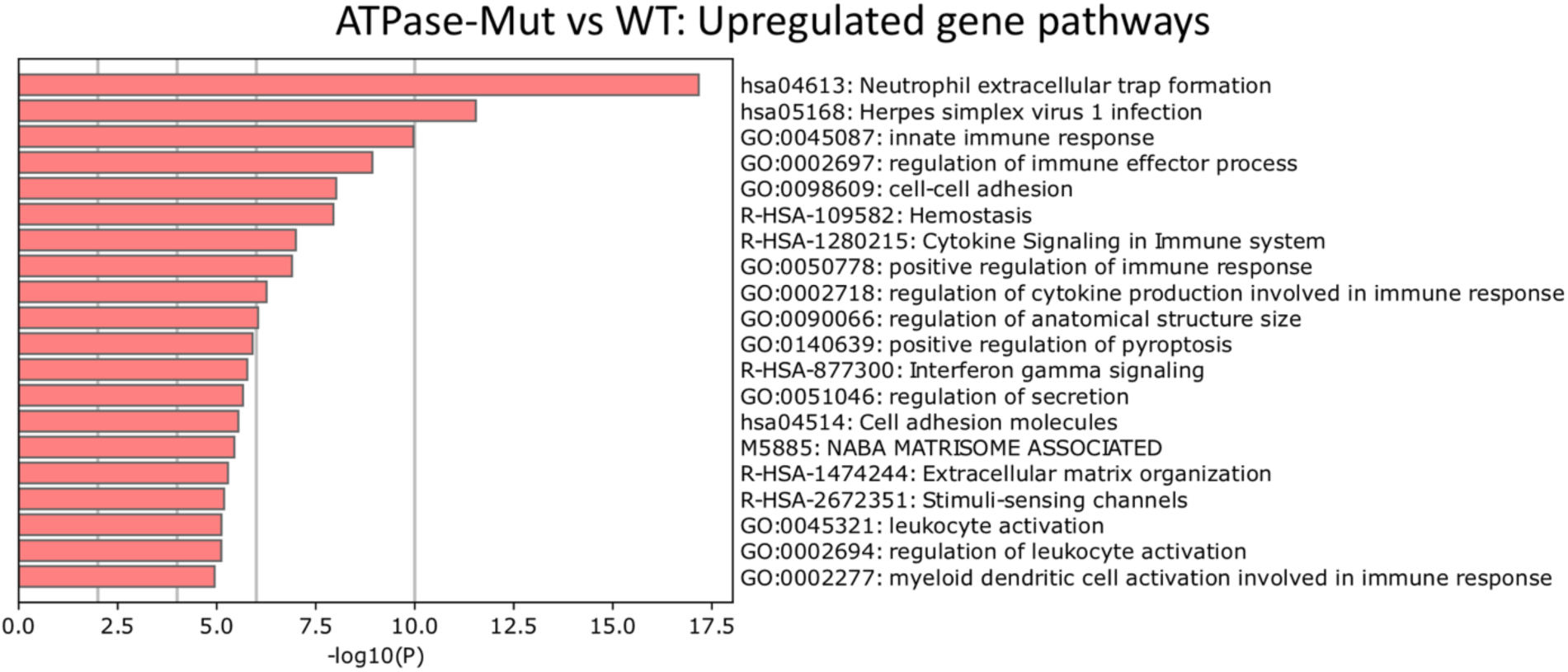
Pathway enrichment for upregulated genes in ATPase-mut vs WT. Pathway enrichment of upregulated genes identified in Figure 4B. Metascape (Zhou et al., 2019) was used for analysis based on the GO Biological Processes, Reactome Pathway, and KEGG Pathway databases. The top 20 pathways are shown.

**Figure S5.**
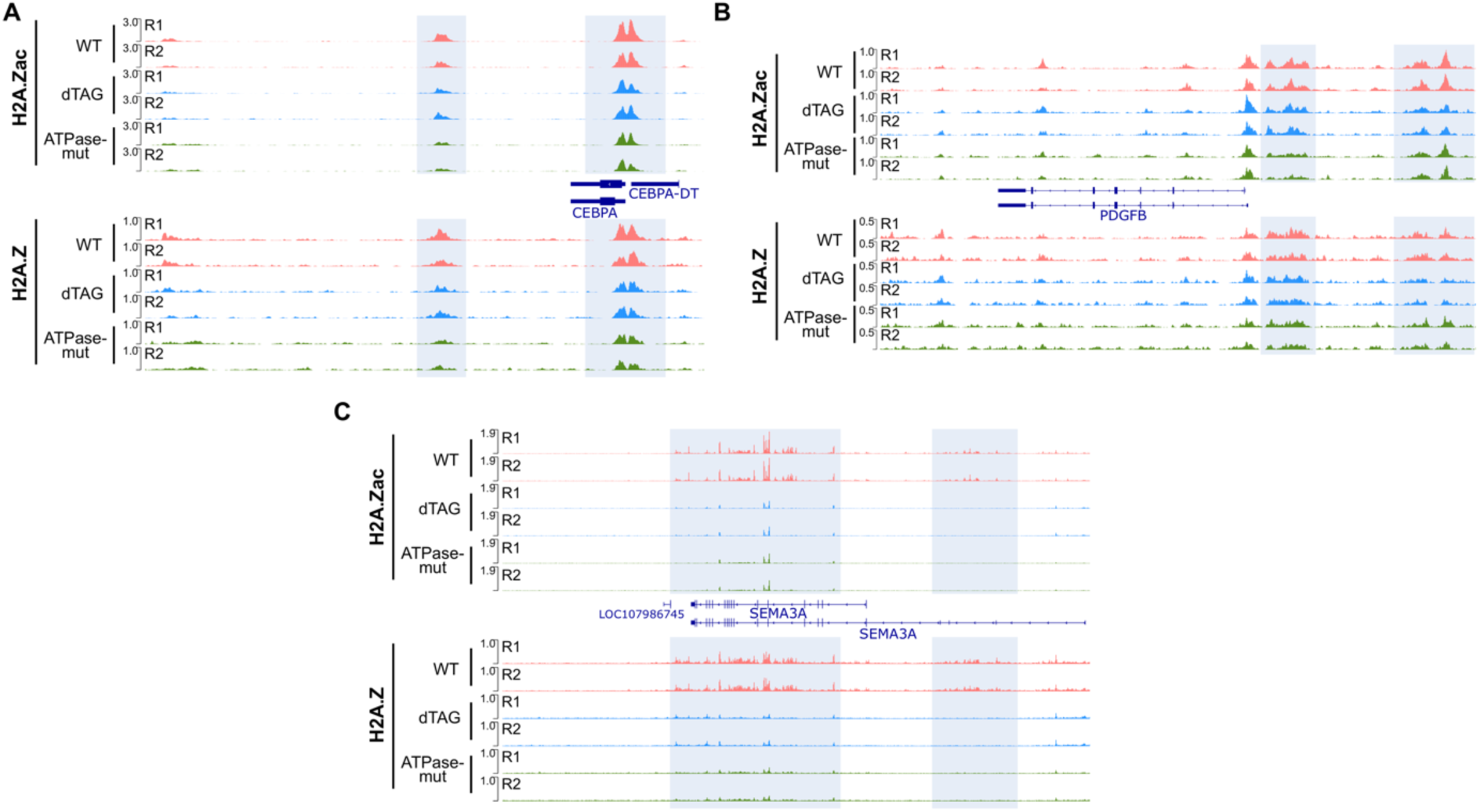
NuA4/TIP60-mediated H2A.Z(ac) deposition at gene promoters regulates transcription. Related to Figure 5. IGV snapshots of RPM-normalized profiles of H2A.Z and H2A.Zac in WT, dTAG and ATPase-mut conditions at promoters of genes downregulated in ATPase-mut vs WT RNA-seq, **(A)** CEBPA, **(B)** PDGFB and **(C)** SEMA3A.

**Figure S6.**
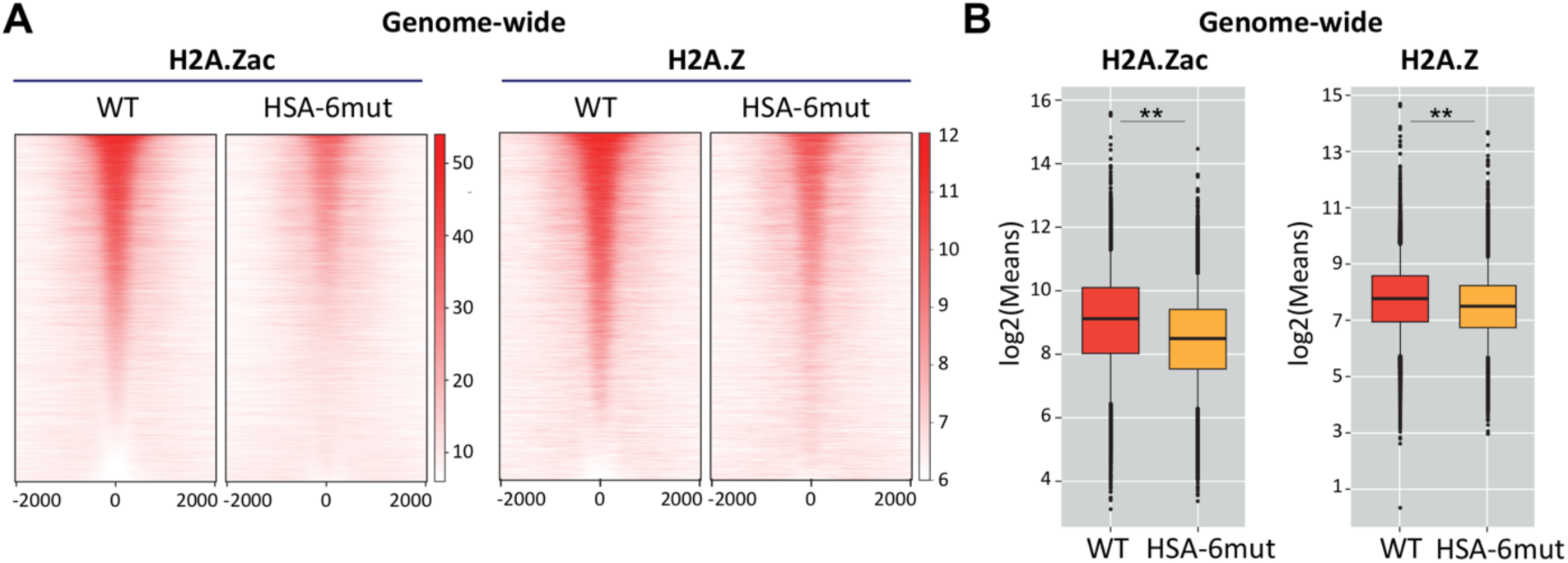
NuA4/TIP60 HAT module is required for H2A.Z remodeling. **(A)** Left, Heatmap representation of genome-wide H2A.Zac peaks which significantly change in HSA-6mut relative to WT, ranked in descending order based on signal in the WT condition. Right, heatmap representation of H2A.Z signal in WT and HSA-6mut specifically for peaks which overlap the H2A.Zac peaks depicted on the left panels. **(B)** Boxplot representation of the heatmaps in (A). ChIP-seq data are merged from two biological replicates (n=2). The read per million (RPM) signals were plotted on the heatmaps. Wilcoxon rank-sum test was performed on the boxplots (\*\**P* < 0.01).

**Figure S7.**
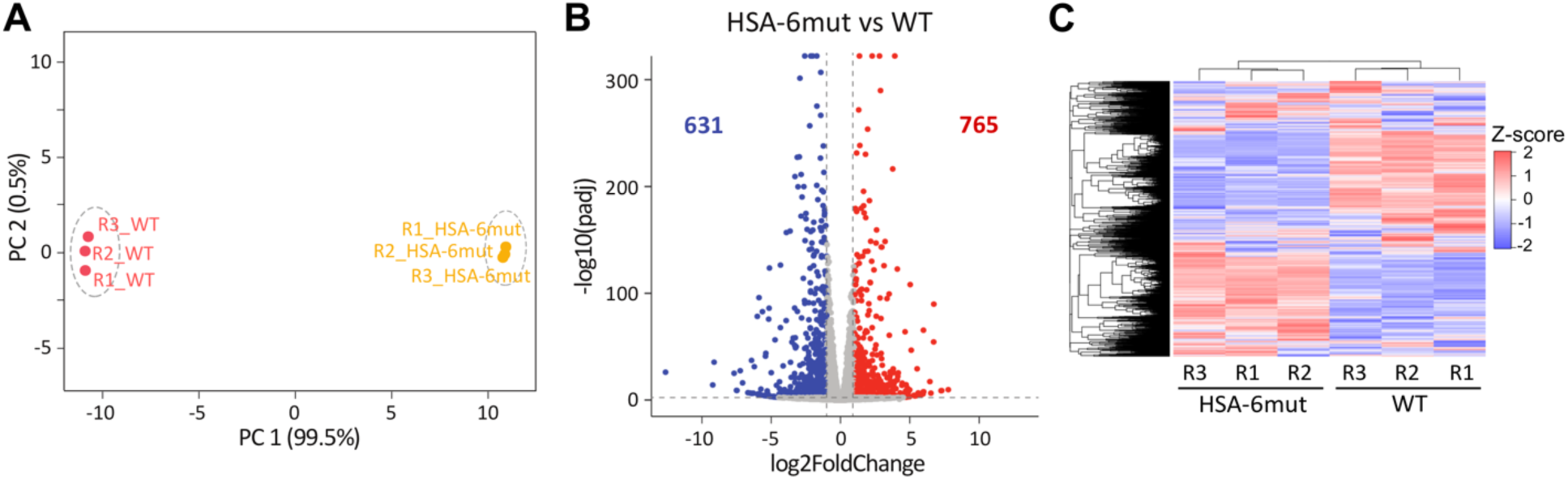
Transcriptional changes resulting from loss of the HAT module. **(A)** PCA plot for WT-complemented and HSA-6mut-complemented RNA-seq samples. **(B)** Volcano plot representation of differential gene expression, with downregulated genes in blue (log2(FC) < -1, padj < 0.01), upregulated genes in red (log2(FC) > 1, padj < 0.01), and unchanged genes in gray. Analysis was performed on three biological replicates (n=3) per condition. **(C)** Hierarchically clustered Z-score heatmap representing normalized transcript counts for HSA-6mut vs WT differentially expressed genes.

**Figure S8.**
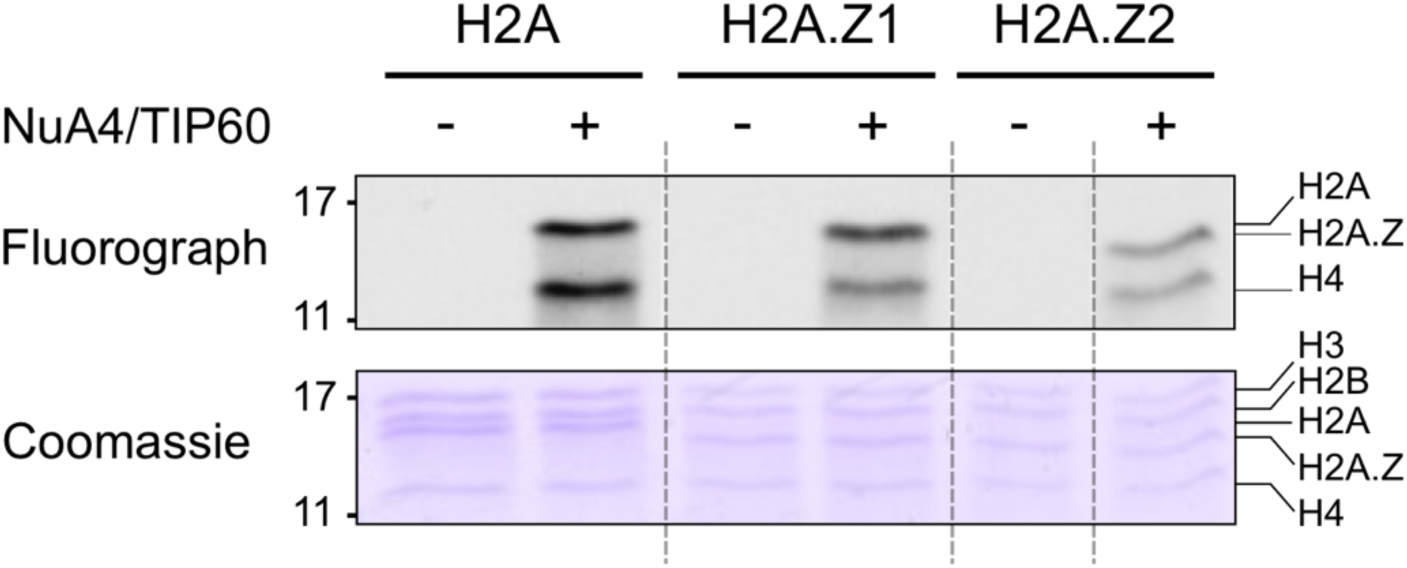
NuA4/TIP60 acetylates nucleosomal H2A.Z. Top, gel fluorograph after radioactive HAT assay using a tandem affinity purified NuA4/TIP60 fraction and different recombinant mononucleosomes as substrates. The mononucleosomes either have H2A (canonical), H2A.Z.1 or H2A.Z2. Bottom, Coomassie staining showing all fours histones in each nucleosome.

